# Rescuing cognitive deficits in alcohol-dependent mice by targeting physiological adaptations in prefrontal cortex K_V_7 channels

**DOI:** 10.1101/2025.11.07.687180

**Authors:** Kathy L. Lindquist, Sudarat Nimitvilai-Roberts, Hyland C. Gonzalez, Jessica H. Hartman, John J. Woodward, Larry S. Zweifel, Jennifer A. Rinker, Patrick J. Mulholland

## Abstract

Chronic alcohol misuse causes cognitive deficits and functional adaptations in the prefrontal cortex (PFC) that facilitate excessive drinking and increase relapse probability during prolonged withdrawal. Positive K_V_7 channel modulation reduces alcohol intake in high-drinking rodents, but the impact of alcohol dependence on cortical K_V_7 physiology and its role in cognitive impairments are understudied. Here, we demonstrate spatial working memory deficits and physiological adaptations in intrinsic excitability and K_V_7 channel function in intratelencephalic PFC projection neurons in alcohol-dependent mice. CRISPR-SaCas9 deletion of *Kcnq3* in PFC neurons that project to the dorsomedial striatum mimicked physiological adaptations and working memory deficits produced by alcohol dependence. Furthermore, PFC microinfusion of the K_V_7 positive modulator retigabine rescued alcohol-induced working memory dysfunction. These findings identify an aberrant cortical mechanism responsible for alcohol-associated cognitive dysfunction, providing insights into a pharmacological treatment approach that can target both high drinking and cognitive impairments associated with alcohol use disorder.

## Introduction

Alcohol use disorder (AUD) is an escalating public health issue that is associated with a general decline in overall health and the emergence of cognitive dysfunction^1–4^. Executive function deficits, such as diminished working memory capacity, are common following alcohol cessation and can lead to an increased probability of relapse^3,5^. The dorsomedial prefrontal cortex (dmPFC) is a nexus of the brain’s executive cognitive system. In humans, alcohol dependence decreases activation of the dmPFC during spatial working memory^1,6,7^, alters dmPFC coherence during cognitive performance^8^, and is associated with volumetric decrease within the prefrontal cortex^1,9^. The rodent dmPFC is also heavily implicated in cognitive function and is susceptible to neuroadaptations produced by alcohol dependence^1,10–12^. Because current pharmacological treatments for AUD are not designed to mitigate cognitive dysfunction, the identification of novel therapeutics that can specifically target cognitive impairments remains necessary to reduce the high rate of relapse in AUD individuals.

Voltage-gated K_V_7 channels are ubiquitously expressed throughout the CNS where they are essential for regulating membrane potential and neuronal excitability^13–15^. K_V_7 channels are arranged as heterotetramers principally composed of K_V_7.2 (*Kcnq2*) and K_V_7.3 (*Kcnq3*) subunits. Within the prefrontal cortex, K_V_7 channels are expressed in all neuronal subtypes^16,17^ and influence cognitive function^13,18–20^. Deep-layer dmPFC pyramidal neurons have diverse projection targets that can be subdivided into intratelencephalic (IT) and extratelencephalic (ET) neurons, distinguished by the presence of *I*_h_ ‘sag’ current^21–23^. IT neurons project contralaterally within the cortex and to other telencephalic regions, such as striatum and amygdala^21,23^. In contrast, ET neurons project to regions outside of the telencephalon, including thalamus and hypothalamus^21,23^. Importantly, IT and ET cortical neurons play distinct roles in cognitive domains. For example, ET neurons display robust activity surrounding choices in an auditory-based decision task^24^, and inhibition of cortical IT, but not ET neurons causes working memory deficits^25^. Acute alcohol (i.e., ethanol) reduces K_V_7 channel currents^26–28^, and preclinical studies have identified K_V_7 channels as modulators of alcohol-related behaviors, including high alcohol drinking^29–31^, alcohol withdrawal-induced negative affect and hyperalgesia^32,33^, and alcohol tolerance^26^. One study showed that concurrent administration of the K_V_7 channel positive modulator retigabine with alcohol for 3 weeks improves cognitive performance in male rats^34^. How alcohol dependence alters K_V_7 channel function within IT and ET cortical projection subtypes to induce dysfunction of cognitive processes and if targeting K_V_7 channels during withdrawal improves spatial working memory remain unknown.

To determine the role of K_V_7 channels in regulating alcohol-induced cognitive impairment, a series of behavioral and electrophysiological studies were performed using the chronic intermittent ethanol (CIE) model of alcohol dependence in male and female C57BL/6J mice. After identifying working memory deficits in CIE-exposed mice, whole-cell patch clamp electrophysiology recordings were performed to measure K_V_7-mediated physiological adaptations in deep-layer dmPFC projection neurons. CRISPR-SaCas9-deletion of *Kcnq3* in dmPFC→dorsomedial striatum (DMS) circuit mimicked CIE-induced physiological changes in dmPFC projection neurons and cognitive dysfunction. Finally, to pharmacologically rescue the physiological deficits, retigabine was microinfused into the dmPFC prior to assessment of working memory.

## Methods

### Mice

All procedures were performed in male and female C57BL/6J (Jackson Laboratory, ME; strain # 000664) mice that were ∼8-10 weeks old at experimental onset. All mice were individually housed within a temperature and humidity-controlled vivarium. Mice were given *ad libitum* access to food and water (Harlan Teklad Diet 2918) and kept on a 12-h reverse light/dark cycle (lights off at 11 am). Animal care was approved by the Medical University of South Carolina Institutional Animal Care and Use Committee in accordance with NIH guidelines for humane care of laboratory animals.

### Delayed Alternation Spatial Working Memory Task

The T-maze protocol was adapted from studies showing working memory deficits in mice exposed to a chronic alcohol liquid diet model^35^. For 5 days, mice were habituated to cupped handling to mitigate handling induced stress during subsequent testing. For two 10-minute sessions on consecutive days, mice were allowed to freely explore the T-maze. On the following day, mice were tested to determine if they successfully choose between alternate arms on a T-maze following a 30-second inter-trial interval (ITI). For each trial, they were confined in their choice arm for 30 seconds before being placed back in the stem arm for 30 seconds to begin the next trial. On the test day, mice underwent seven of these trials, followed by an 8^th^ trial with a 5-second ITI to ensure that mice remained motivated to complete the task. Any urine or fecal matter produced during ongoing trials was removed with water to avoid becoming a visual or odor cue. Mice underwent this 3-day process before beginning CIE to establish a baseline working memory level. Based on their initial performance, mice were pseudo-randomly assigned to control or CIE treatment groups. The task was then repeated 72-96 h following the final CIE session to assess cognitive deficits with ITIs of 30 or 90 seconds. Performance on the working memory task was reported as percent correct alternations.

### Alcohol dependence model

To establish alcohol dependence, mice were exposed to the CIE model^36^. Mice were injected with the alcohol dehydrogenase inhibitor pyrazole (1 mmol/kg; IP; Sigma–Aldrich, St. Louis, MO), dissolved in 1.6 g/kg ethanol (8% w/v in 0.9% v/v sterile saline; IP; 20 ml/kg dose volume), before placement in plexiglass ethanol inhalation chambers (Plas Labs, Inc, Lansing, MI) for a 16-h exposure period, followed by an 8-h withdrawal period. Ethanol concentration within the chambers was monitored so that blood ethanol levels were ∼ 200-250 mg/dl. BECs were measured via retro-orbital blood samples (40 μL) collected with heparinized capillary tubes (Drummond Scientific Company, Broomall, PA) after each session. After samples were centrifuged at 10000×g for 10 minutes, plasma was separated for BEC measurement using an alcohol analyzer (Analox Instrument, Stourbridge, UK). Control mice were exposed to air and injected with pyrazole dissolved in saline, such that all mice received the same number of injections and handling. CIE vapor was administered Monday through Thursday, followed by a 72-h withdrawal period. This cycle was repeated four times, as this amount of CIE exposure has been shown to cause cognitive deficits when tested 72 h after the final CIE session^11^.

### Whole-cell patch-clamp slice electrophysiology

Mice were anesthetized via isoflurane before rapid decapitation. Coronal brain sections including the dmPFC (i.e., prelimbic and anterior cingulate cortex) were collected using a Leica VT 1200S vibratome before placement into oxygenated artificial cerebral spinal fluid (ACSF) and incubation at 34 C for 30 minutes, followed by 30 minutes at room temperature. Slices were transferred to the recording chamber and continually perfused with oxygenated ACSF. Cutting ACSF solution during slice preparation was composed of the following: 2.5 mM KCl, 1.25 mM NaH_2_PO_4_, 0.4 mM C_6_H_8_O_6_, 10 mM C_6_H_12_O_6_, 200 mM sucrose, 25 mM CHNaO_3_, 6 mM MgCl_2_, 0.5 mM CaCl_2_. Recording ACSF solution was composed of the following: 125 mM NaCl, 2.5 mM KCl, 1.25 mM NaH_2_PO_4_, 0.4 mM C_6_H_8_O_6_, 10 mM C_6_H_12_O_6_, 25 mM CHNaO_3_, 1.3 mM MgCl_2_, 2 mM CaCl_2_, 2.4 mM sodium pyruvate. Reagents for ACSF and internal solutions were purchased from Sigma Aldrich. Recordings were conducted using Axon Multiclamp 700B amplifier and the Instrutech ITC 18 analog digital converter, which were controlled by Axograph software. Thin wall borosilicate glass electrodes (Sutter instruments) were pulled on a Sutter Instrument P97 micropipette puller with tip resistances ranging from 2-3.5 MΩ. Internal solution for recordings where evoked action potential firing and M-current were collected was composed of the following: 10 mM HEPES, 0.1 mM EGTA, 4 mM ATP-Mg, 2 mM NaGTP, 10 mM Na-phosphocreatine salt hydrate, 110 mM KCl, 10 mM tetrapotassium pyrophosphate. Internal solution for evoked action potential firing for recording from dmPFC →DMS projection neurons was composed of the following: 10 mM HEPES, 1mM EGTA, 2 mM NaATP, 0.3 mM NaGTP, 2 mM MgCl_2_, 10 mM KCl, and 120 mM K-gluconate. Neurons whose resistance changed by over 20% during the recordings were excluded from analysis.

Deep-layer pyramidal neurons were identified within the dmPFC of mice based on location and neuronal morphology. To identify IT vs ET neurons, a 200-pA hyperpolarizing current was applied to generate *I*_h_ sag current. This allowed for the calculation of the physiological index (PI) using the following equation^37^:

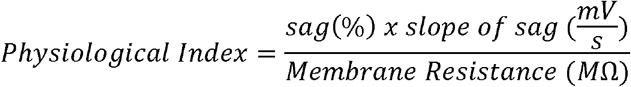

Recordings were acquired in current clamp mode to measure evoked action potential firing, which was induced with a series of direct current injections in increments of 20 pA for 1 s from 0-300 pA. All recordings were analyzed for total action potentials generated at each current step, cumulative action potential generation, rheobase, resting membrane potential, action potential half-width, amplitude, rise time, and afterhyperpolarization using the Easy Electrophysiology Software.

M-current was recorded in voltage clamp in the presence of 200 nM of tetrodotoxin (Alomone Labs, Jerusalem, Israel) following established protocols^14,38^. In whole cell mode, neurons were initially held at −70 mV, and series resistance was compensated at 80%. A depolarizing protocol consisting of five voltage steps incrementing in 10 mV was applied at −70 mV. Following this baseline recording, the K_V_7 positive modulator retigabine (30 μM, Alomone Labs) was bath applied before repeating the depolarizing protocol. Lastly, the K_V_7 antagonist XE-991 (20 μM, Tocris Bioscience, Bristol, UK) was bath-applied to abolish M-current. Using Axograph software offline, the final recording in the presence of XE-991 was subtracted from the initial recording to yield the K_V_7 channel-specific current.

### Stereotaxic surgery

Mice were anesthetized using isoflurane and head-fixed into a stereotaxic instrument (David Kopf Instruments, Tujunga, CA) for the duration of the surgical procedure. An incision was performed to expose the skull, the following coordinates were used to target the dmPFC: A/P +1.78 mm, M/L ±0.4 mm, and D/V −2.25 mm (relative to Bregma). Microinjection cannulas (Protech International, Boerne, TX) targeting both hemispheres were chronically implanted into the dmPFC. Mice underwent postoperative care for a minimum of 5 days following surgery. Retigabine (10 ng/hemisphere; dihydrochloride, Alomone Labs) was dissolved in sterile 0.9% saline and microinfused 30 minutes before T-maze testing at a dose volume of 0.15 μl/min over 2 min. Cannula placements were verified using alcian blue staining. Mice were excluded if the cannula tip was outside of the dmPFC.

The following coordinates were used for infusion of CRISPR-SaCas9 *Kcnq3* vectors: dmPFC: A/P +1.78 mm, M/L ±0.4 mm, and D/V −2.25 mm (relative to Bregma), DMS: A/P +1.05 mm, M/L ±1.50 mm, and D/V −3.0 mm. To specifically knockdown *Kcnq3* expression within dmPFC→DMS projection neurons, a retrograde Cre virus (AAVrg-Ef1a-mCherry-IRES-Cre, #55632-AAVrg, Addgene) was infused into the DMS (300 nL) and a Cre-dependent CRISPR construct was infused into the dmPFC (200 nL; control: AAV1-FLEX-SaCas9-U6-sgRosa26 + AAV1-FLEX-EYFP, *Kcnq3* knockdown: AAV1-FLEX-SaCas9-U6-sgKcnq3 + AAV1-FLEX-EYFP, viral vectors courtesy of Dr. Zweifel). For validation of protein, 200 nl of either CRISPR vector was infused into the dmPFC with AAV8-Ef1a-mCherry-IRES-Cre (#55632-AAV8, Addgene) and AAV1-FLEX-EYFP. All mice underwent postoperative care for a minimum of 5 d following surgery. For CRISPR studies, mice used for behavioral testing were sacrificed for histological confirmation following task completion 4 weeks after initial baseline testing.

### Brain tissue punch homogenization

One-millimeter brain tissue punches were collected from the dmPFC of each mouse and homogenized in 50 µL of 2% SDS buffer (Hoefer, Inc., Bridgewater, MA, USA) supplemented with protease and phosphatase inhibitors (Thermo Fisher Scientific, Waltham, MA, USA). Tissue punches were placed in Axygen microcentrifuge tubes (Corning, Corning, NY, USA) containing approximately 25 µL of 0.5-mm zirconia/silica beads (BioSpec Products, Bartlesville, OK, USA). Samples were homogenized in a Bullet Blender Storm Pro (Next Advance, Troy, NY, USA) at 66.7% power for 30 seconds, followed by a 1-minute incubation on ice. This cycle was repeated four times. Homogenates were transferred to clean 1.5-mL microcentrifuge tubes and centrifuged at 20000 × g for 10 minutes at 4°C. The supernatant was transferred to a new tube, and total protein concentration was determined using the Pierce BCA Protein Assay Kit (Thermo Fisher Scientific). The remaining supernatant and pellet were aliquoted and stored at −80°C until further analysis.

### Quantification of K_V_7.3 protein

K_V_7.3 protein abundance was quantified using the Jess Simple Western automated western blot platform (ProteinSimple, Minneapolis, MN, USA). Samples were thawed on ice and diluted to a final protein concentration of 1.0 mg/mL in 0.1× sample buffer prepared from 10× Sample Buffer 2 supplied with the 12–230 kDa Separation Module (ProteinSimple). Fivefold master mix, prepared using EZ Standard Pack 1 (ProteinSimple), was added at one-fifth of the final sample volume. Three samples with starting protein concentrations below 1.0 mg/mL were not further diluted before addition of the master mix. Samples were vortexed, denatured at 95°C for 5 minutes, vortexed again, and briefly centrifuged. Samples were analyzed using a Jess Simple Western system and a 25-well assay plate. K_V_7.3 was detected using a rabbit anti-K_V_7.3 primary antibody diluted 1:10 in Antibody Diluent 2 (Synaptic Systems, Göttingen, Germany), followed by a ready-to-use anti-rabbit secondary antibody (ProteinSimple). Total protein was measured using the Total Protein Detection Module (ProteinSimple) and used to normalize K_V_7.3 abundance.

### Physiological and behavioral validation of CRISPR-SaCas9-deletion of Kcnq3

Mice were sacrificed ∼4-5 months following survival surgery for slice electrophysiology. dmPFC neurons projecting to the DMS were identified based on EYFP fluorescence. Recordings were acquired in current clamp mode to measure evoked action potential firing and biophysical membrane properties as described above. For working memory assessment, mice underwent habituation and pre-deletion testing on the T-maze task starting 5 d after surgery. Mice were then retested 4 weeks after pre-deletion testing when deletion is persistent^39^. Due to perseverative responses during baseline testing, two control mice were removed from analysis.

### Statistics

T-tests, multi-factorial ANOVA for between-subjects designs, or mixed-effects models for repeated-measures designs were used for analyses with Bonferroni post hoc tests to correct for multiple comparisons, when appropriate (GraphPad Prism 10.2.3). For the 5 s ITI data, correct and incorrect alternations were classified as binary responses and analyzed using a quasibinomial GLM in R (R-project.org). Working memory performance in Figure 1 was analyzed using a 4-factor analysis in R. An α of 0.05 was set as a threshold for statistical significance. Sex was included as a full factor in all analyses. All data are reported mean ± SEM.

**Fig 1.**
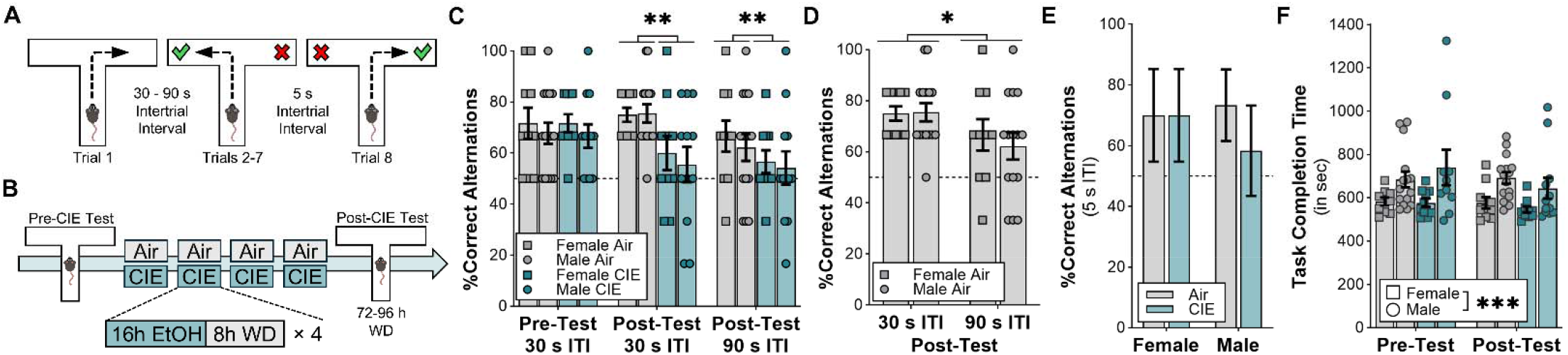
Chronic intermittent ethanol exposure caused spatial working memory deficits in male and female C57BLZ6J mice. **(A)** Schematic of delayed alternation task with 2 different **ITIs** for trials 2-7 and a 5 s ITI for the 8^th^ trial. (B) Timeline for testing working memory performance before and after CIE exposure. **(C)** Quantification of correct trial percentage on the delayed alternation task for the 30 s ITI and 90 s ITI versions of the task between CIE exposed and control male and female mice (4-factor RM analysis, Treatment x Time interaction: *F*_1,137_ = 4.069, *p* = 0.045; n = 10-15 mice/group/sex; post hoc tests: Air, post-CIE vs. CIE, post-CIE, *p* = 0.0025; CIE pre vs. post: *p* = 0.039). **(D)** Quantification of correct trial percentage between the 30 s ITI and 90 s ITI for Air control male and female mice (2-factor RM analysis, ITI main effect: *F*_1,46_ = 5.887, *p* = 0.019). **(E)** Comparison of correct alternations on the 5 s ITI between CIE exposed and control mice (quasibinomial GLM, p = −0.68 ± 0.865, *t=* −0.781, *p* = 0.439). **(F)** Comparison of total time to complete the task between CIE and control mice pre- and post-CIE exposure (3-factor RM analysis, Treatment main effect: *F*_143_ = 0.026, *p* = 0.876; Sex main effect: *F*_1,43_ = 14.72, *p* = 0.0004).

## Results

### Chronic alcohol-induced spatial working memory deficits

Previous reports show that the CIE exposure model induces deficits in some cognitive domains during withdrawal^11,40–44^. Here, a delayed alternation task was used to determine if spatial working memory deficits would emerge in alcohol-dependent mice (**Fig. 1A,B**). Performance was assessed prior to and after CIE exposure, and ITI delays of 30 and 90 s were used for the post-CIE tests. During baseline, working memory performance in male and female mice was 70.1 ± 1.19%. There was a significant decrease in working memory performance in the alcohol-dependent mice at both ITIs (**Fig 1C**). In air controls, performance was significantly worse during ITIs of 90 vs. 30 s (**Fig 1D**), demonstrating impaired performance as cognitive load increased. However, all mice exhibited equal spatial working memory on the 8^th^ trial with the 5 s ITI regardless of their treatment history (**Fig 1E**). To determine if the deficit emerged due to locomotor effects, as some have reported that C57BL/6J mice can be hypoactive during early withdrawal^45^, the total time required to complete the delayed alternation task was quantified. While CIE did not impact task completion time (**Fig. 1F**), females performed the task significantly faster than males regardless of treatment (**Fig. 1F**), suggesting that female and male mice use different cognitive strategies, as sex hormones can influence spatial-based task strategies^46,47^.

### dmPFC physiological adaptations in alcohol-dependent mice

To determine a possible mechanism underlying CIE-induced spatial working memory impairment, intrinsic excitability was measured in deep-layer dmPFC pyramidal neurons. At 72-96 h withdrawal from CIE, whole-cell patch-clamp electrophysiological recordings were performed in layer V dmPFC pyramidal neurons (**Fig. 2A**). Analysis of evoked firing revealed a significant increase in evoked action potentials in neurons from alcohol-dependent male and female mice (**Fig. 2A,B**). CIE exposure also significantly increased the total number of evoked action potentials fired (**Fig. 2C**), cumulative action potentials fired (**Supplementary Fig. 1a**), decreased rheobase (**Fig. 2D**), and depolarized the resting membrane potential (**Fig. 2E**) with no changes to action potential kinetics (**Supplemental Table 1**; *p* values > 0.05).

**Fig 2.**
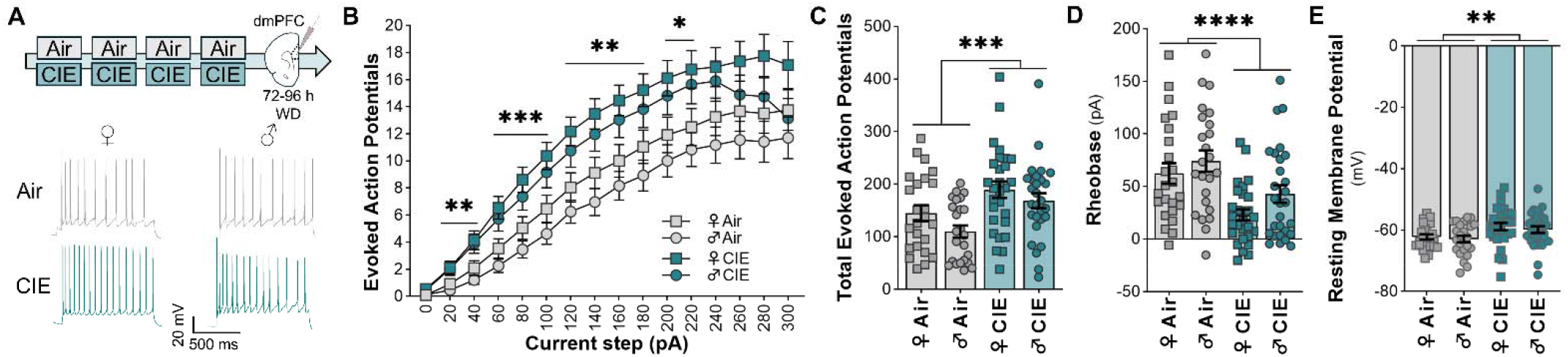
Alcohol dependence increased neuronal excitability of layer V dmPFC neurons. **(A)** Experimental timeline of alcohol exposure and whole-cell patch-clamp electrophysiology targeting layer V dmPFC pyramidal neurons with representative traces of evoked action potentials. **(B)** Line plot displaying the number of action potentials fired between 0-300 pA at 20 pA intervals in Air control and CIE-exposed mice (3-factor analysis, Voltage Step x Treatment interaction: *F*_2.26,230.6_ = 3.922, *p* = 0.017; *n = 7-8* mice/sex/treatment). **(C)** Quantification of total evoked action potentials across summed across all current steps (2-factor analysis, Treatment main effect: *F*_1,100_ = 12.88, *p* = 0.0005). **(D)** Measurement of rheobase in Air control and CIE-exposed mice (2-factor analysis, Treatment main effect: *F*_1,100_ = 17.77, *p* = 0.00006). **(E)** Comparison of resting membrane potential between Air control and CIE exposed mice. (2-factor analysis, Treatment main effect: *F*_1,100_ = 9.139, *p* = 0.0032).

Because K_V_7 channels control neural firing, the resting membrane potential, and cognition^13,14,20,48–50^, we next determine if CIE-induced neuronal excitability adaptations were related to a decrease in K_V_7 channel function. K_V_7-specific current (i.e., M-current) was recorded using a standard depolarizing voltage step protocol^13,14,38^. Following baseline recording, the current was first modulated using bath application of retigabine (30 μM) and then blocked by bath application of the K_V_7 blocker XE-991 (20 μM). Analysis showed a significant reduction in M-current amplitude at −60 mV (*p* = 0.0066) and −20 mV (*p* = 0.0307) voltage steps in CIE-exposed mice (**Fig. 3A-B**). Because M-current was markedly larger at the most depolarized voltage command, the −20-mV step was analyzed separately. At this voltage step, M-current significantly differed by sex, which revealed larger M-current in dmPFC neurons from female mice (**Fig. 3C**), and CIE exposure significantly reduced M-current amplitude in both sexes (**Fig. 3C**). For retigabine-modulated M-current, there was a significant interaction of Voltage Step × Treatment (**Fig. 3D-E**), although post hoc comparisons did not reach statistical significance (**Fig. 3D-E**; *p* > 0.06). When the −20-mV voltage step was analyzed separately, M-current amplitude after retigabine bath application was significantly lower in dmPFC neurons from alcohol-dependent mice (**Fig. 3F**). Thus, M-current and retigabine-modulation of M-current were reduced by alcohol dependence in layer V dmPFC projection neurons.

**Fig 3.**
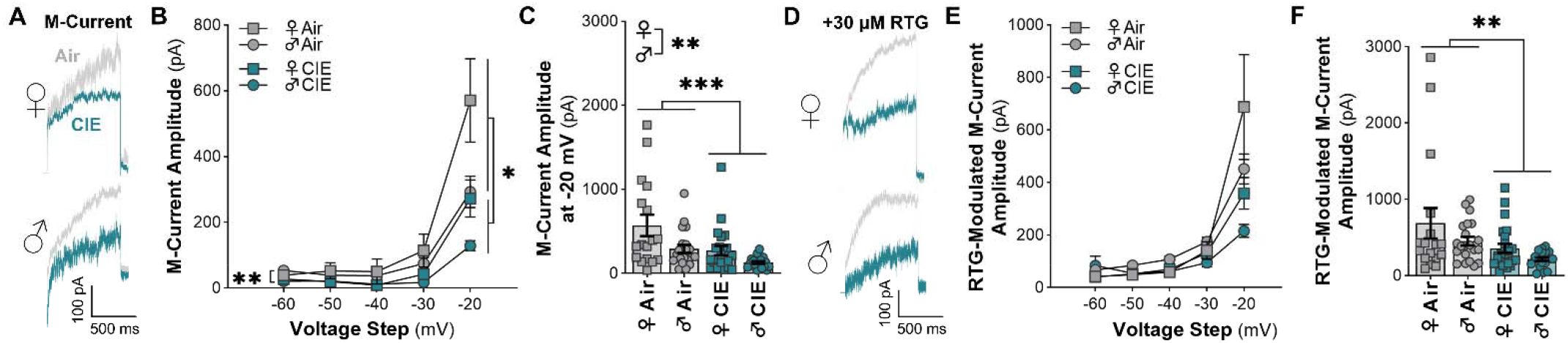
K_v_7-mediated M-current amplitude is reduced in layer V dmPFC projection neurons recorded from alcohol-dependent mice. **(A)** Representative M-current traces. **(B)** M-current amplitude across increasingly depolarized voltage steps in Air control and CIE-exposed mice (3-factor RM analysis, Voltage Step x Treatment interaction, *F*_1.183,92.30_ = 5.895, *p* = 0.013; *n = 7-8* mice/sex/treatment). **(C)** Comparison M-current amplitude at the −20 mV voltage step (2-factor analysis, Sex main effect, *F*_1,76_ = 9.532, *p* = 0.0028; Treatment main effect, *F*_1,76_ = 11.44, *p* = 0.0011). **(D)** Representative M-current traces when slices were bath applied with the K_v_7 channel positive modulator retigabine. **(E)** M-current responses following retigabine wash in from −60 mV-+20 mV in 10 mV steps (3-factor RM analysis, Voltage Step x Treatment interaction, *F*_1.129.88,88.10_ = 6.310, *p* = 0.011). **(F)** Comparison of M-current amplitude in the presence of retigabine for the −20 mV voltage step (2-factor analysis, Treatment main effect, *F*_1,76_ = 8.390, *p* = 0.0049).

### Differentiating CIE effects on ET and IT projection neurons

Deep-layer IT and ET pyramidal neurons have diverse projection targets and distinct physiological properties^21–23^. Thus, recordings from control mice shown in **Fig. 2** were categorized into ET and IT neurons based on potential physiological differences in sag ratio (**Fig. 4A,B**) and membrane resistance (**Fig. 4C**), and a physiological index was calculated to separate the two distinct populations of projection neurons. When graphed based on their sag ratio and membrane resistance, IT and ET neurons largely form their own distinct clusters (**Fig. 4D**), and analysis showed that ET neurons have a significantly larger physiological index in both female and male dmPFC projection neurons (**Fig. 4E**). Next, we determined the influence of CIE exposure on intrinsic excitability and M-current amplitude in these two projection neuron subpopulations.

**Fig 4.**
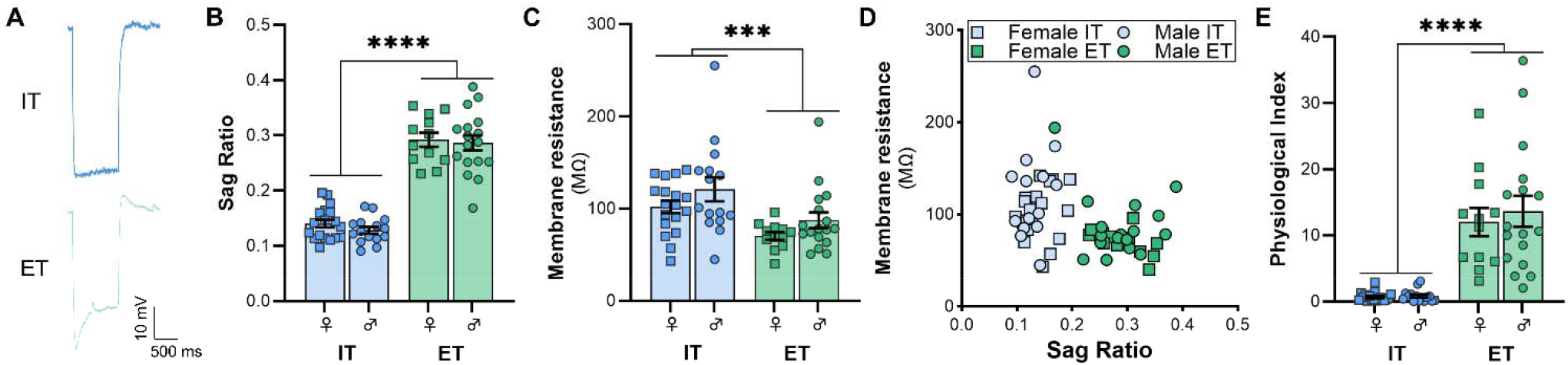
Subtypes of layer V dmPFC projection neurons can be separated based on electrophysiological properties. **(A)** Representative sag current traces from IT and ET neurons. **(B)** Calculation of sag ratio between IT and ET projection neurons (2-factor analysis, projection main effect: *F*_1,58_ = 223.4, *p* < 0.000001). **(C)** Comparison of membrane resistance between IT and ET neurons (2-factor analysis, Projection main effect: *F*_1,58_ = 12.89, *p* = 0.0007). **(D)** Scatter plot demonstrating sag ratio and membrane resistance in IT and ET neurons. (E) Quantification of physiological index for IT and ET neurons (2-factor analysis, Projection main effect: *F*_1,58_ = 60.54, *p* < 0.000001).

Within the ET population, analysis of evoked firing revealed a significant Treatment × Current Step × Sex interaction (**Fig. 5A,B**). While follow-up 2-factor interactions were not significant, there was a significant main effect of Treatment showing an increase in evoked firing in alcohol-dependent mice (**Fig. 5B**). CIE exposure did not affect total evoked action potential firing (**Fig. 5C**), although there was a significant reduction in rheobase (**Fig. 5D**) and a more depolarized resting membrane potential (**Fig. 5E**). CIE exposure did not affect other action potential kinetic properties (**Supplemental Table 2**), and there was a modest effect of CIE on cumulative spiking, but only at one current step (**Supplementary Fig. 1b**). Next, M-current within the ET population was measured (**Fig. 5F,G**), and there was no effect of CIE exposure on amplitude across all voltage steps (**Fig. 5G**). When analyzing M-current at the −20 mV step, M-current amplitude was larger in female dmPFC ET neurons (**Fig. 5H**) but was not changed by CIE (**Fig. 5H**). Retigabine modulated M-current did not differ by CIE exposure across all voltage steps (**Fig. 5J**) or when the −20 mV step was analyzed separately (**Fig. 5K**). Based on these findings, ET neurons were largely insensitive to physiological changes induced by CIE exposure, except for a potential increase in intrinsic excitability in male dmPFC ET projection neurons and subtle yet significant changes in the RMP and rheobase.

**Fig 5.**
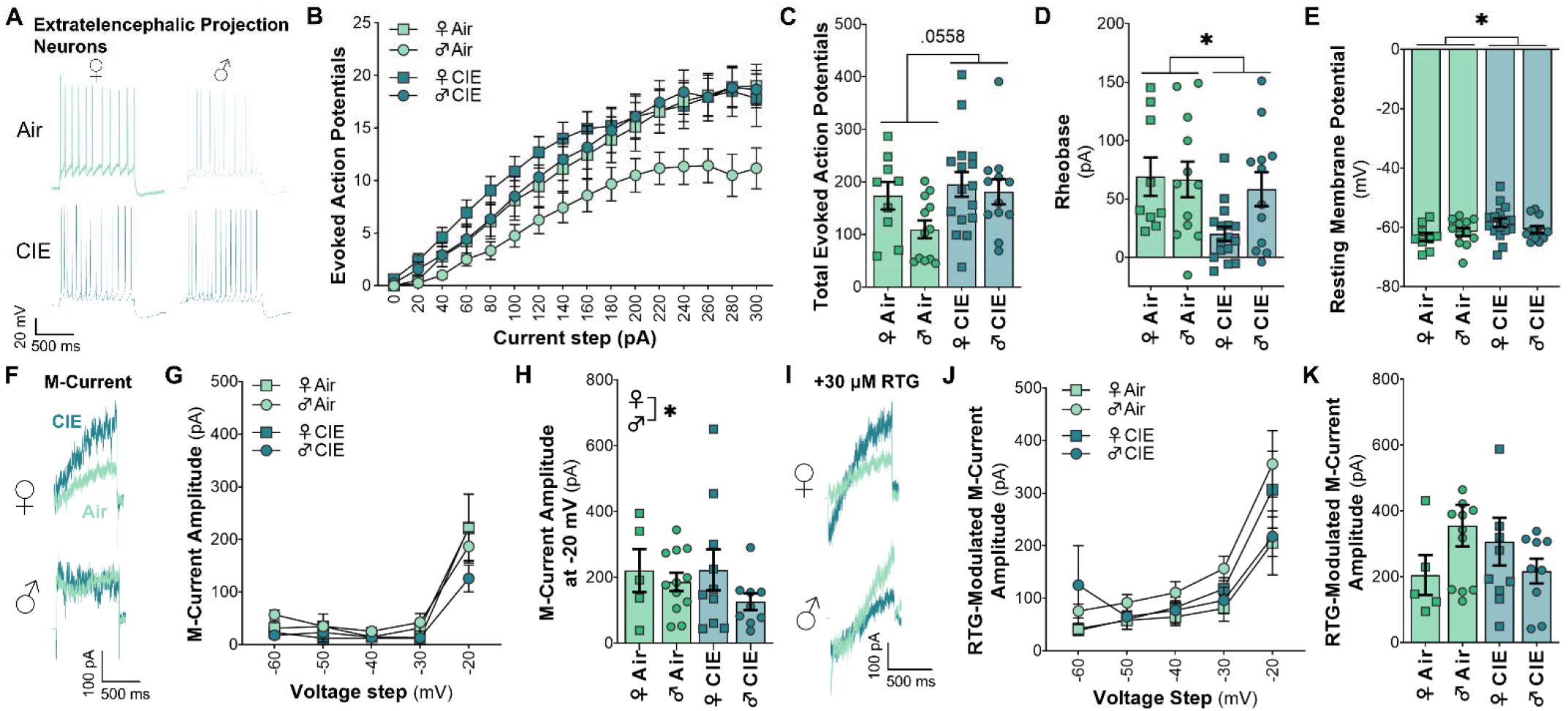
Layer V dmPFC ET projection neurons showed subtle excitability adaptations without changes in M-current. **(A)** Representative traces of evoked action potentials in ET neurons. **(B)** Action potentials generated in ET projection neurons across 0-300 pA current steps in 20 pA increments (3-factor RM analysis, Treatment x Current Step x Sex interaction: *F*_2.387,107.4_ = 3.393, *p* < 0.0295; Treatment main effect: *F*_1,47_ = 4.789, *p* = 0.0336). **(C)** Summation of action potentials generated across all current step applied to ET neurons (2-factor analysis, Treatment main effect: *F*_1,45_ = 3.855, *p* = 0.0558). **(D)** Comparison of rheobase between ET neurons from Air control and CIE-exposed mice (2-factor analysis, Treatment main effect: *F*_1,45_ = 4.984, *p* = 0.0306). **(E)** Measurement of resting membrane potential (2-factor analysis, Treatment main effect: *F*_1,45_ = 4.132, *p* = 0.0480). **(F)** Representative M-current traces in ET neurons. **(G)** Measurement of M-current amplitude generated from multiple voltage steps (3-factor RM analysis, Treatment main effect: *F*_1,32_ = 2.785, *p* = 0.1049). **(H)** Comparison of M-current amplitude at the −20 mV voltage step across treatment groups (3-factor RM analysis, Sex main effect: *F*_1,33_ = 4.380, *p* = 0.0441; Treatment main effect: *F*_1,33_ = 2.111, *p* = 0.1557). **(I)** Representative M-current traces from ET neurons following bath application of retigabine. **(J)** Retigabine modulated M-current across all voltage steps (3-factor RM analysis, Treatment main effect: *F*_1,160_ = 3.203, *p* = 0.0754). **(K)** Comparison of M-current amplitude at the −20 mV voltage step in the presence of bath application of retigabine (2-factor analysis, Treatment main effect: *F*_1,32_ = 0.074, *p* = 0.7874).

Within the IT dmPFC population, CIE exposure significantly increased evoked firing (**Fig. 6A,B**). Total evoked action potentials in IT neurons were also significantly increased in male and female alcohol-dependent mice (**Fig. 6C**), as was cumulative action potentials fired (**Supplementary Fig. 1c**). CIE-exposed IT neurons had a significantly lower rheobase (**Fig. 6D**) and depolarized resting membrane potential (**Fig. 6E**). Other action potential kinetic properties are shown in **Supplemental Table 3**. Upon analyzing M-current in IT neurons (**Fig 6F**), there was a significant Treatment × Voltage Step interaction, and post hoc comparisons showed decreased M-current at the −20 mV step in neurons from alcohol-dependent mice (**Fig. 6G**). There were also main effects of Treatment (**Fig. 6H**) and Sex (**Fig. 6H**) at the −20 mV step, where CIE reduced M-current amplitude and M-current was larger in female neurons compared to males. There was a significant Treatment × Voltage Step interaction in retigabine modulated M-current (**Fig. 6I-J**), and post hoc comparisons revealed a significant difference between Air and CIE at the −20 mV step (*p* = 0.014). Further analysis of the maximal step confirmed a diminished retigabine modulated M-current amplitude in neurons from CIE-treated mice (**Fig. 6K**). When considering the overall physiological adaptations in dmPFC neurons from alcohol-dependent mice, these results demonstrate that the changes in intrinsic excitability and K_V_7 channel function are largely driven by adaptations in IT projection neurons. Interestingly, M-current was significantly larger in IT neurons compared with ET neurons from Air control mice (−20 mV step: *F*_1,34_ = 9.463, *p* = 0.0041; **Fig 5G vs. 6G**).

**Fig 6.**
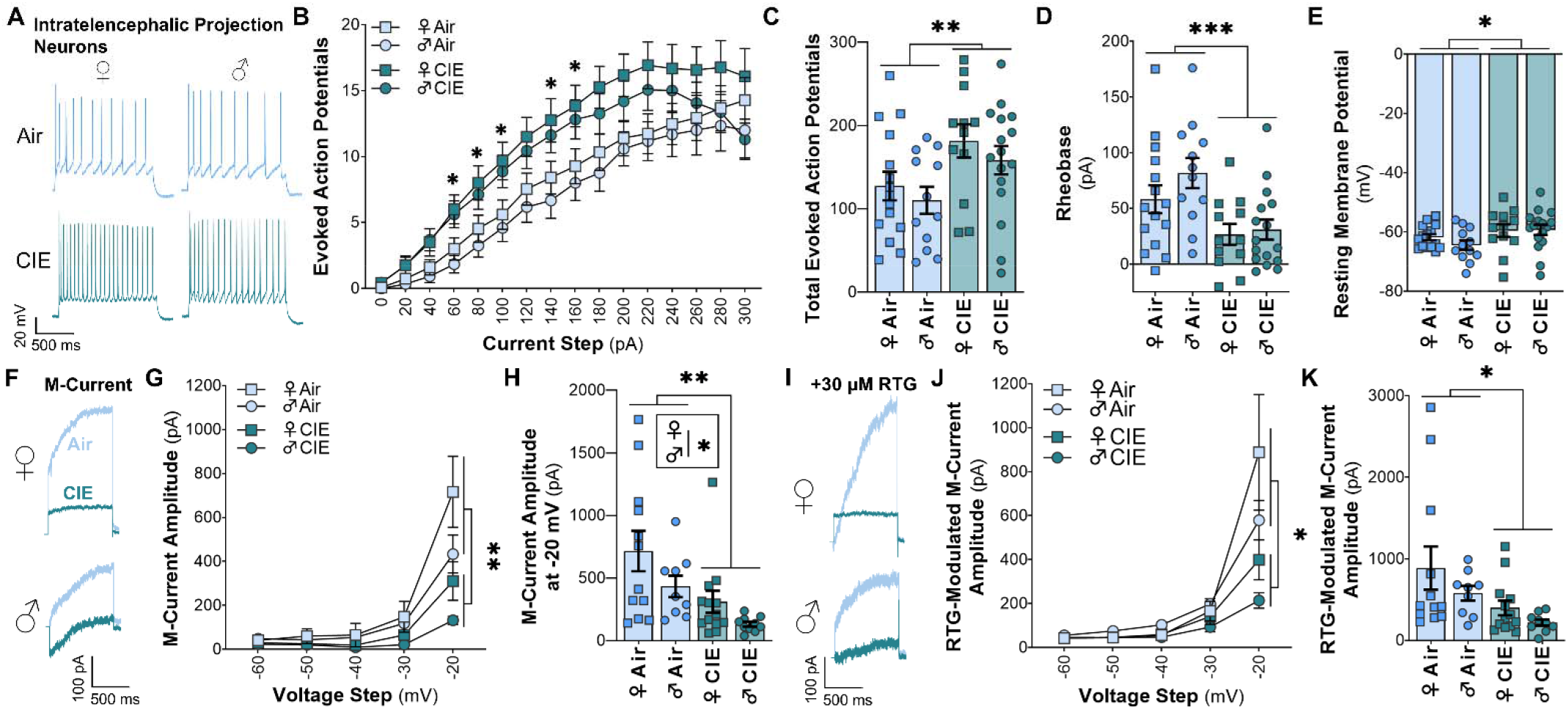
Alcohol dependence increased neuronal excitability and reduced M-current in layer V dmPFC IT neurons. **(A)** Representative action potential traces from IT neurons. **(B)** Action potentials generated in IT projection neurons across 0-300 pA current steps in 20 pA increments (3-factor RM analysis, Treatment x Current Step interaction: *F*_2.321,132.3_ = 3.223, *p* = 0.0359, post hoc tests: 60 - 160 pA: *p’s* < 0.0427). **(C)** Summation of action potentials generated across all current step applied to IT neurons (2-factor analysis, Treatment main effect: *F*_1,55_ = 7.134, *p* = 0.0099). **(D)** Comparison of rheobase between IT neurons from Air control and CIE-exposed mice (2-factor analysis, Treatment main effect: *F*_1,51_ = 13.26, *p* = 0.0006). **(E)** Measurement of resting membrane potential (2-factor analysis, Treatment main effect: *F*_1,51_ = 4.944, *p* = 0.031). **(F)** Representative M-current traces in IT neurons. **(G)** Measurement of M-current amplitude in IT neurons generated from multiple voltage steps (3-factor RM analysis, Treatment x Voltage Step interaction: *F*_1.190,49.97_ = 7.775, *p* = 0.0051, −20 mV: *p* = 0.0036). **(H)** Comparison of M-current amplitude at the −20 mV voltage step across treatment groups (2-factor analysis, Treatment main effect: *F*_1,40_ = 10.40, *p* = 0.0025; Sex main effect: *F*_1,40_ = 4.419, *p* = 0.0419). **(I)** Representative M-current traces from IT neurons following positive modulation with retigabine. **(J)** Retigabine modulated M-current across all voltage steps (3-factor RM analysis, Treatment x Voltage Step interaction: *F*_1.058,44.43_ = 7.017, *p* = 0.0101; post hoc, *p* = 0.014). **(K)** Comparison of M-current amplitude at the −20 mV voltage step in the presence of bath application of retigabine (2-factor analysis, Treatment main effect: *F*_1,40_ = 7.082, *p* = 0.0112).

### Kcnq3 deletion in dmPFC→DMS projection neurons mimicked CIE-induced physiological changes and cognitive deficits

IT projection neurons in the dmPFC send dense axonal projections to the DMS where they are known to control working memory performance^51^. We hypothesize that reduced M-current in IT neurons that project to the DMS contributes to CIE-induced spatial working memory deficits. A previously characterized CRISPR construct to knock down *Kcnq3*^39,52^ was used to confirm a causal role of K_V_7 channels in dmPFC→DMS circuitry for cognitive control over working memory performance. Thus, deletion of *Kcnq3* in this circuit should mimic physiological changes in dmPFC projection neurons and spatial working memory deficits in alcohol-dependent male and female mice. For validation of K_V_7.3 protein knockdown, mice receive co-injections of AAV1-FLEX-SaCas9-U6-sg*Kcnq3* and AAV1-FLEX-EYFP (to visualize cells) with AAV8-Ef1a-mCherry-IRES-Cre into the dmPFC, and control mice received injections of AAV1-FLEX-SaCas9-U6-sgRosa26 in place of the sg*Kcnq3* vector. As expected, CRISPR-deletion of *Kcnq3* significantly reduced K_V_7.3 protein expression in dmPFC (**Fig. 7A,B**). Spatial working memory was then assessed 5 d post-surgery during the viral transduction and early genome editing (pre-deletion) phase then again after 4 weeks during the persistent *Kcnq3* knockdown phase^39^. Correct performance during the pre-deletion test averaged 76.1 ± 8.41 and 72.2 ± 4.81 in Rosa26 and sg*Kcnq3* mice, respectively. Spatial working memory did not differ in Rosa26 control mice across time (Unpaired t-test: *t*_5_ = 0.707, *p* = 0.256). However, CRISPR-deletion of *Kcnq3* in dmPFC→DMS projection neurons (**Fig. 7C,D**) significantly reduced working memory when compared to their own pre-deletion testing performance (**Fig. 7E**) and to the Rosa26 control mice during the post-deletion test (**Fig. 7F**). Patch-clamp recordings showed that CRISPR-deletion of *Kcnq3* significantly increased dmPFC→DMS projection neuron evoked firing (**Fig. 7G,H**), including increases in total (**Fig. 7I**) and cumulative (**Supplementary Fig. 1d**) action potentials fired. *Kcnq3* deletion also reduced rheobase in dmPFC projection neurons (**Fig. 7J**). Notably, EYFP expression was absent in dmPFC in mice that did not receive AAVrg infusion into the DMS (data not shown). Together, these results demonstrate that CRISPR-deletion of *Kcnq3* in dmPFC→DMS projection neurons mimicked behavioral deficits and physiological changes in IT neurons during withdrawal from CIE exposure.

**Fig 7.**
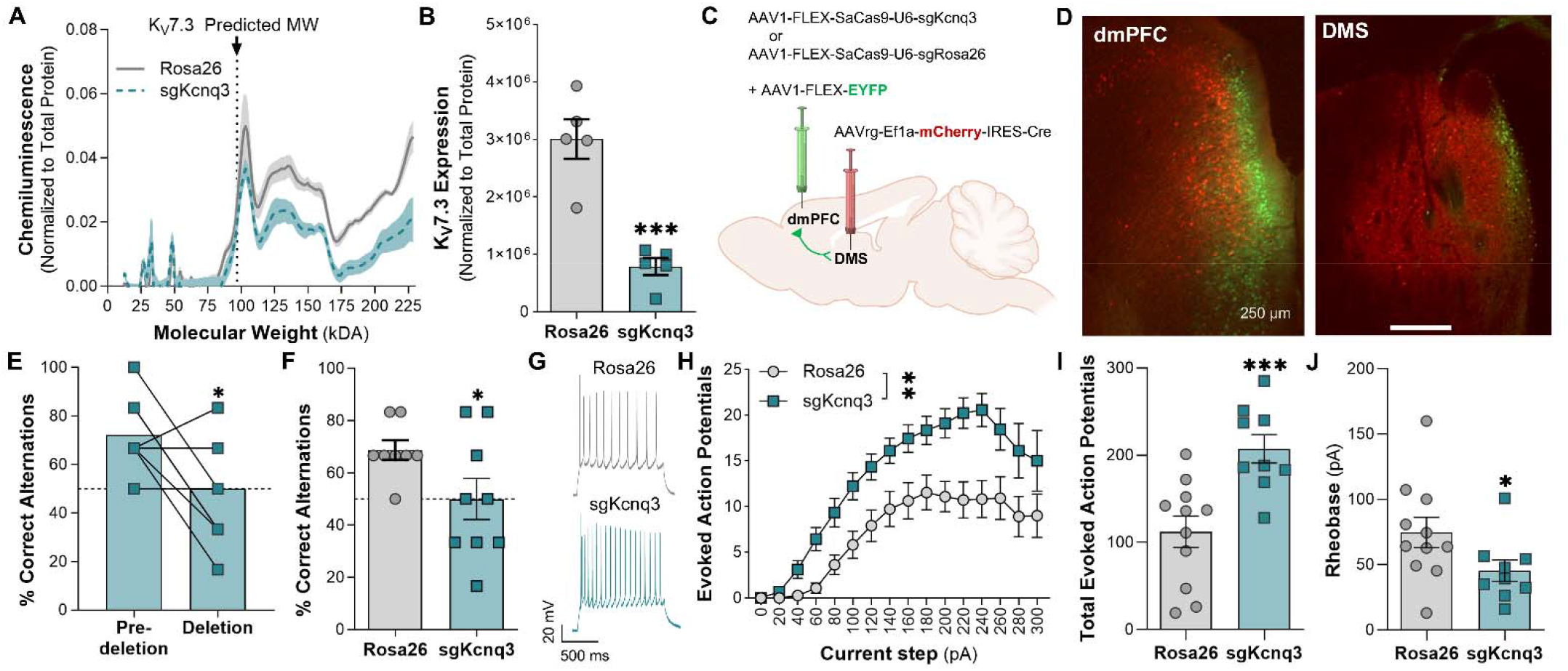
CRISPR-SaCas9 deletion of *Kcnq3* in dmPFC^DMS projection neurons mimicked alcohol dependence-induced hyperexcitability and spatial working memory dysfunction. **(A)** Quantification of chemiluminescence following deletion of *Kcnq3* within the dmPFC. **(B)** Comparison of K_v_7.3 protein expression between control and CRISPR-SaCas9 deletion of *Kcnq3* within the dmPFC (Unpaired t-test: *t_8_* = 5.898, *p* = 0.0004; *n* = 5 mice/group). **(C)** Diagram of viral approach for CRISPR-SaCas9 deletion of *Kcnq3* in dmPFC—>DMS projection neurons. **(D)** Representative images of mCherry and EYFP expression in the dmPFC (left panel) and DMS (right panel). **(E)** Comparison of delayed alternation performance pre- and post­deletion of *Kcnq3* in the dmPFC->DMS circuit (Paired t-test: *t_8_* = 2.219, *p* = 0.0286; *n* = 8-9 mice/vector). **(F)** Comparison of delayed alternation performance between *Kcnq3* deletion and sgRosa26 controls at the post­deletion time point (Unpaired t-test: *t*_15_ = 2.064, *p* = 0.0284). **(G)** Representative action potential traces from Rosa26 and sgKcnq3 neurons. **(H)** Action potentials generated from 0-300 pA at 20 pA increments within the dmPFC—>DMS circuit (2-factor analysis, Vector main effect: *F*_1,18_ = 14.72, *p* = 0.0012; *n* = 2-3 mice/vector). **(I)** Calculation of total evoked action potentials across all current step in dmPFC-^DMS projection neurons (Unpaired t-test: *t*_18_ = 3.836, *p* = 0.0006). **(J)** Comparison of rheobase between control and *Kcnq3* knockdown (Unpaired t-test: *t*_18_ = 1.972, *p* = 0.0321).

### Positive modulation of dmPFC K_V_7 channels rescued cognitive function

Because physiological characterization identified a mechanistic change in K_V_7 channel function, pharmacological modulation of dmPFC K_V_7 channels was performed to determine if retigabine microinfusion would restore spatial working memory performance in alcohol-dependent mice. Spatial working memory was assessed during baseline and after CIE exposure in mice with guide cannula implanted in dmPFC (**Fig. 8A**). Prior to CIE exposure, baseline performance did not differ across groups (Male Air: 78.28 ± 3.93%, Female Air: 74.08 ± 4.04%, Male CIE: 74.08 ± 4.04%, Female CIE: 75.70 ± 4.78%). As expected, CIE exposure significantly reduced working memory performance compared to air controls in male and female mice administered vehicle (**Fig. 8B**). Retigabine microinfusion into the dmPFC of alcohol-dependent mice significantly increased spatial working memory performance compared to CIE mice that received vehicle (**Fig. 8B**). In air controls, performance did not differ between saline and retigabine microinfusion (post hoc: *p* = 0.999). Cannula placements for the location of microinfusions are shown in **Fig. 8C**. These results demonstrate that positive modulation of dmPFC K_V_7 channels rescued cognitive function in alcohol-dependent male and female mice.

**Fig 8.**
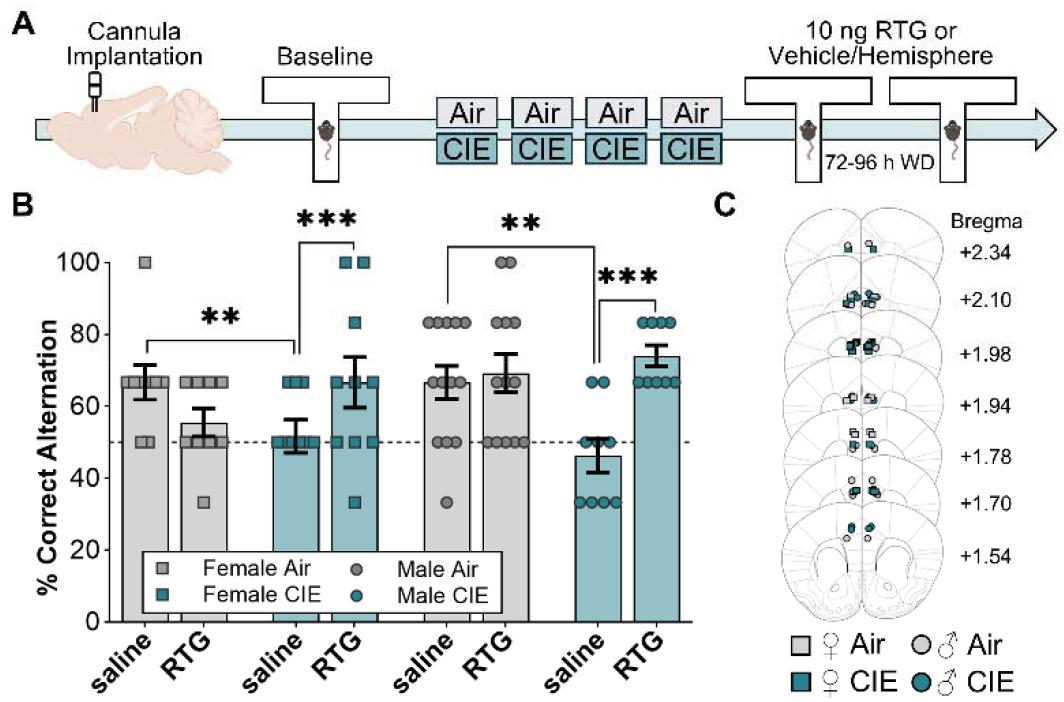
Positive modulation of dmPFC K_v_7 channels rescued spatial working memory in CIE-exposed male and female mice. **(A)** Timeline for microinjection study. **(B)** Quantification of performance in the delayed alternation task in control and CIE-exposed mice microinfused with either saline or retigabine into the dmPFC (3-factor RM analysis, Treatment x Drug interaction, *F*_1,78_ = 11.31, *p* = 0.0012; post hoc: CIE, saline vs. retigabine, *p* = 0.0003; saline, Air control vs. CIE, *p* = 0.0017; *n* = 9-13 mice/treatment/sex). **(C)** Diagram of guide cannula microinjection locations.

## Discussion

Consistent with dysfunction in other cognitive domains^11,40–44^, we found that the CIE model of dependence caused spatial working memory deficits in both male and female mice. In parallel to these behavioral deficits, alcohol dependence increased the excitability of layer V dmPFC projection neurons while concurrently reducing K_V_7-mediated M-current specifically in IT projection neurons. CRISPR-SaCas9-deletion of K_V_7.3 channels in dmPFC→DMS neurons mimicked physiological adaptations and cognitive deficits observed in alcohol-dependent mice, and positive modulation of dmPFC K_V_7 channels with retigabine rescued cognitive function in both sexes of alcohol dependent mice. Together, these findings demonstrate circuit-specific adaptations in cortical K_V_7 channels as a driver of dependence-induced cognitive impairment.

One major finding from these studies is that dmPFC IT, but not ET projection neurons were sensitive to dependence-induced physiological changes in cell firing and M-current. While M-current was larger in female IT neurons, mice from both sexes showed a CIE-induced reduction in M-current amplitude. The mechanisms for IT neuron sensitivity to alcohol are unclear, but we showed that M-current in control mice was more prominent in IT neurons. Although K_V_7 channels are voltage-dependent, they require plasma membrane phosphatidylinositol 4,5-bisphosphate (PI(4,5)P_2_) for stabilization of the ion channel pore to allow the flow of potassium ions, and K_V_7 channels are suppressed by G_q_-coupled M1 and M3 muscarinic receptor activation^13–15^. To our knowledge, no evidence exists for differential expression of K_V_7 channels or PI(4,5)P_2_ across the two subpopulations of dmPFC projection neurons, but acute alcohol can diminish expression of PI(4,5)P_2_ and alter sensitivity of K_V_7 channels to PI(4,5)P_2_^19,27,28^. Thus, the reported effects of CIE may be caused by lower abundance of K_V_7 channels in ET neurons or altered PI(4,5)P_2_ activity specifically in IT projection neurons. As K_V_7 channels are modulated by muscarinic receptors, it is also possible that CIE exposure may disrupt muscarinic modulation of M-current in IT dmPFC projection neurons. It is also known that cholinergic signaling in PFC neurons is projection-specific in that ET neurons show larger and longer excitatory responses after muscarinic activation^53^. Alcohol exposure during adolescence blunts PFC acetylcholine release in rats^54^, and CIE exposure in adult mice decreases acetylcholine release and the density of cholinergic interneurons in the DMS^55^. Future studies should determine how alcohol dependence alters acetylcholine release or muscarinic cholinergic receptor expression in dmPFC subpopulations of projection neurons, to assess whether potential differences could contribute to the increased sensitivity of IT neurons.

Another major finding from these studies is that CIE-induced deficits in spatial working memory were rescued following direct positive modulation of K_V_7 channels within the dmPFC. Although studies show that blocking K_V_7 channels enhances cognitive performance^18,20,48,49,56^, genetic mutation in K_V_7 channels that diminish K_V_7 function can cause cognitive deficits^50,57,58^. This suggests that K_V_7 inhibition operates on an inverted U-shaped curve to regulate cognition. Indeed, Galvin et al^49^ demonstrated an inverted U-shaped curve when modulating working memory in nonhuman primates using a muscarinic agonist, whose actions were directly attributed to K_V_7 channels. In *Drosophila*, KCNQ is known to modulate acute alcohol’s interference with aversive associative memory formation^58^, demonstrating cross-species K_V_7 channel control of alcohol-associated memory impairment. While retigabine has been shown to prevent alcohol-induced cognitive deficits in rats^34^, retigabine was administered concurrently with alcohol in that study. Here, we extended these findings linking K_V_7 specifically to spatial working memory by demonstrating complete rescue of cognitive performance by a single administration of retigabine into the dmPFC during withdrawal. As acute blockade of K_V_7 channels in neurotypical conditions improves cognitive performance, increasing channel activity in neuropsychiatric conditions associated with reduced cortical K_V_7 function can improve cognition. Thus, these findings demonstrate that K_V_7 channels, while sensitive to changes induced by alcohol dependence, are a viable treatment target to restore proper cognitive performance.

Our physiological recordings demonstrated that IT projection neurons were sensitive to alcohol dependence and positive modulation of K_V_7 channels within the dmPFC rescued the cognitive deficits. However, the specific projections driving impairments in spatial working memory in alcohol dependent mice are unknown. The brain regions involved in spatial working memory include prefrontal cortex, hippocampus, mediodorsal thalamus, and nucleus reuniens^59–62^, all of which are sensitive to CIE-induced physiological adaptations^11,63–65^. However, a study demonstrated that PFC neurons that project to the DMS also control working memory performance^51^. Given the reduction in M-current specifically in IT projection neurons, we reasoned that reduced K_V_7 channel currents in dmPFC neurons that project to the DMS drive working memory deficits in alcohol-dependent mice. Indeed, CRISPR-SaCas9 deletion of *Kcnq3* in dmPFC→DMS projection neurons revealed K_V_7 channel control over spatial working memory in this circuit. In demonstrating this, these studies also revealed that selective deletion of K_V_7 channels in dmPFC→DMS projections mimicked the physiological changes in neural excitability and spatial working memory behavioral deficits observed in CIE-exposed mice. Together, these findings causally implicate down-regulation of K_V_7 channels in dmPFC→DMS projection neurons as a key mechanism of cognitive impairments associated with alcohol dependence.

In summary, the current findings support the effectiveness of K_V_7 channel positive modulation to rescue chronic alcohol-induced cognitive impairment. Together with our previous studies showing that retigabine reduces alcohol drinking preferentially in rodents with high drinking phenotypes^29,30^, these results further highlight the potential for K_V_7 channel positive modulators as an option for treating individuals with AUD. AUD is often comorbid with depression^66^, and recent findings from clinical trials support K_V_7 channel openers as an emerging class of therapeutics to treat major depressive disorder through modulation of reward circuits^67–70^. Thus, K_V_7 channel positive modulators may treat multiple symptoms of AUD, including heavy drinking, cognitive impairment, and co-morbid mood disorders that drive high rates of relapse.

## Acknowledgements

K.L., J.R., and P.M. designed the experiments. K.L., H.G., J.H., S.R., and P.M. performed the experiments and analyzed the data. K.L., J.R., J.W., L.Z., J.H., and P.M. interpreted the data. K.L. and P.M. wrote the paper, and all authors provided editorial input on the manuscript.

K.L.L was supported by NIH award F31 AA031433. P.J.M. was supported by NIH awards P50 AA010761 and R01 AA023288.

## Data availability

Date will be made available upon publication and request.

