## Supplementary Materials for "Rescuing cognitive deficits in alcohol-dependent mice by targeting physiological adaptations in prefrontal cortex K_V_7 channels"

**Supplementary Tables**

**Supplementary Table 1**. Action potential kinetics of deep layer dmPFC pyramidal neurons from Figure 2.

|  | Membrane Resistance (MΩ) | Capacitance (pF) | Spike Time  (s) | Peak  (mV) | Amplitude  (mV) | Threshold  (mV) | Rise Time (ms) | Decay Time (ms) | Half-Width (ms) | fAHP  (mV) | mAHP  (mV) |
| --- | --- | --- | --- | --- | --- | --- | --- | --- | --- | --- | --- |
| ♀  Air | 90.39 ±  6.38 | 26.93 ±  1.27 | 0.41 ±  0.06 | 44.33 ±  1.20 | 84.73 ±  1.70 | -40.39 ±  1.23 | 0.43 ±  0.02 | 2.61 ±  0.10 | 1.88 ±  0.05 | -10.65 ±  1.35 | -15.62 ±  0.84 |
| ♂  Air | 104.60 ±  10.35 | 24.17±  1.21 | 0.35 ±  0.05 | 42.23 ±  1.46 | 81.03 ±  1.86 | -38.80 ±  0.96 | 0.43 ±  0.03 | 2.70 ±  0.15 | 1.91 ±  0.08 | -10.29 ±  2.00 | -16.08 ±  0.67 |
| ♀  CIE | 100.97 ±  8.10 | 23.34 ±  1.10 | 0.39 ±  0.05 | 43.54 ±  1.44 | 85.50 ±  1.60 | -41.84 ±  1.15 | 0.42 ±  0.02 | 2.57 ±  0.12 | 1.93 ±  0.09 | -8.98 ±  2.18 | -15.51 ±  0.78 |
| ♂ CIE | 113.57 ±  11.04 | 23.61 ±  1.14 | 0.38 ±  0.04 | 43.66 ±  1.60 | 83.34 ±  1.40 | -39.79 ±  1.05 | 0.45 ±  0.02 | 2.81 ±  0.14 | 1.94 ±  0.07 | -13.99 ±  1.52 | -16.70 ±  0.74 |

**Supplementary Table 2**. Action potential kinetics of deep layer ET dmPFC pyramidal neurons from Figure 5. Bold denotes main effect of CIE treatment, *p* <0.05.

|  | Membrane Resistance (MΩ) | Capacitance (pF) | Spike Time  (s) | Peak  (mV) | Amplitude  (mV) | Threshold  (mV) | Rise Time (ms) | Decay Time (ms) | Half-Width  (ms) | fAHP  (mV) | mAHP  (mV) |
| --- | --- | --- | --- | --- | --- | --- | --- | --- | --- | --- | --- |
| ♀ Air | 66.31 ±  4.31 | **30.20 ±**  **2.70** | 0.40 ±  0.05 | 45.43 ±  2.00 | 87.77 ±  2.49 | -42.34 ±  2.34 | 0.38 ±  0.03 | 2.27 ±  0.10 | 1.77 ±  0.09 | -9.71 ±  1.57 | -13.35 ±  1.28 |
| ♂ Air | 84.63 ±  11.79 | **26.00 ±**  **1.66** | 0.37 ±  0.08 | 43.14 ±  1.77 | 81.40 ±  2.87 | -38.26 ±  1.38 | 0.39 ±  0.02 | 2.27 ±  0.08 | 1.70 ±  0.04 | -9.28 ±  1.75 | -15.54 ±  0.90 |
| ♀ CIE | 82.39 ±  7.03 | **25.01 ±**  **1.21** | 0.41 ±  0.06 | 44.41 ±  1.61 | 86.37 ±  2.10 | -41.96 ±  1.59 | 0.39 ±  0.02 | 2.25 ±  0.11 | 1.78 ±  0.13 | -6.55 ±  3.57 | -15.20 ±  0.72 |
| ♂ CIE | 82.44 ±  12.30 | **25.62 ±**  **2.14** | 0.37 ±  0.06 | 44.26 ±  2.80 | 85.24 ±  2.47 | -40.98 ±  1.35 | 0.45 ±  0.03 | 2.26 ±  0.16 | 1.74 ±  0.14 | -12.32 ±  1.93 | -14.69 ±  0.84 |

**Supplementary Table 3**. Action potential kinetics of deep layer IT dmPFC pyramidal neurons from Figure 6. Bold denotes main effect of CIE treatment, *p* <0.05.

|  | Membrane Resistance (MΩ) | Capacitance (pF) | Spike Time  (s) | Peak  (mV) | Amplitude  (mV) | Threshold  (mV) | Rise Time (ms) | | Decay Time (ms) | Half-Width  (ms) | fAHP  (mV) | mAHP  (mV) |
| --- | --- | --- | --- | --- | --- | --- | --- | --- | --- | --- | --- | --- |
| ♀ Air | 104.84 ±  7.80 | **24.97 ±**  **1.00** | 0.42 ±  0.09 | 43.67 ±  1.52 | 82.90 ±  2.21 | -39.23 ±  1.36 | 0.46 ±  0.03 | 2.82 ±  0.12 | | 1.95 ±  0.05 | -11.21 ±  1.97 | -17.07 ±  0.97 |
| ♂ Air | 124.58 ±  15.39 | **22.34 ±**  **1.67** | 0.33 ±  0.05 | 41.33 ±  2.37 | 80.65 ±  2.48 | -39.33 ±  1.37 | 0.46 ±  0.05 | 3.13 ±  0.24 | | 2.13 ±  0.13 | -11.30 ±  3.67 | -16.62 ±  0.99 |
| ♀ CIE | 125.74 ±  13.71 | **21.11 ±**  **1.87** | 0.36 ±  0.07 | 42.67 ±  3.13 | 84.34 ±  2.53 | -41.67 ±  1.73 | 0.46 ±  0.04 | 2.99 ±  0.19 | | 2.13 ±  0.07 | -12.23 ±  1.50 | -16.61 ±  1.42 |
| ♂ CIE | 136.92 ±  14.70 | **22.11 ±**  **1.10** | 0.38 ±  0.05 | 43.01 ±  1.46 | 81.91 ±  1.55 | -38.91 ±  1.53 | 0.46 ±  0.02 | 3.23 ±  0.14 | | 2.09 ±  0.05 | -15.23 ±  2.23 | -18.24 ±  0.98 |

**Supplementary Table 4**. Action potential kinetics of dmPFC→DMS neurons from Figure 7. Bold font denotes *p* < 0.05.

|  | Membrane Resistance (MΩ) | Capacitance (pF) | Spike Time  (s) | Peak  (mV) | Amplitude  (mV) | Threshold  (mV) | Rise Time (ms) | | Decay Time (ms) | Half-Width  (ms) | fAHP  (mV) | mAHP  (mV) |
| --- | --- | --- | --- | --- | --- | --- | --- | --- | --- | --- | --- | --- |
| sgRosa | 74.46 ±  8.08 | 13.32 ±  1.98 | 0.35 ±  0.03 | 48.35 ±  3.99 | 81.08 ±  5.10 | -32.73 ±  2.09 | 0.62 ±  0.06 | 2.14 ±  0.04 | | 2.55 ±  0.17 | **-15.32 ±**  **1.60** | -10.78 ±  1.57 |
| sgKcnq3 | 70.07 ±  6.42 | 17.84 ±  2.66 | 0.42 ±  0.06 | 41.26 ±  5.36 | 78.40 ±  5.21 | -37.14 ±  1.26 | 0.63 ±  0.08 | 2.06 ±  0.04 | | 2.46 ±  0.22 | **-8.54**  **± 2.80** | -10.63 ±  0.90 |

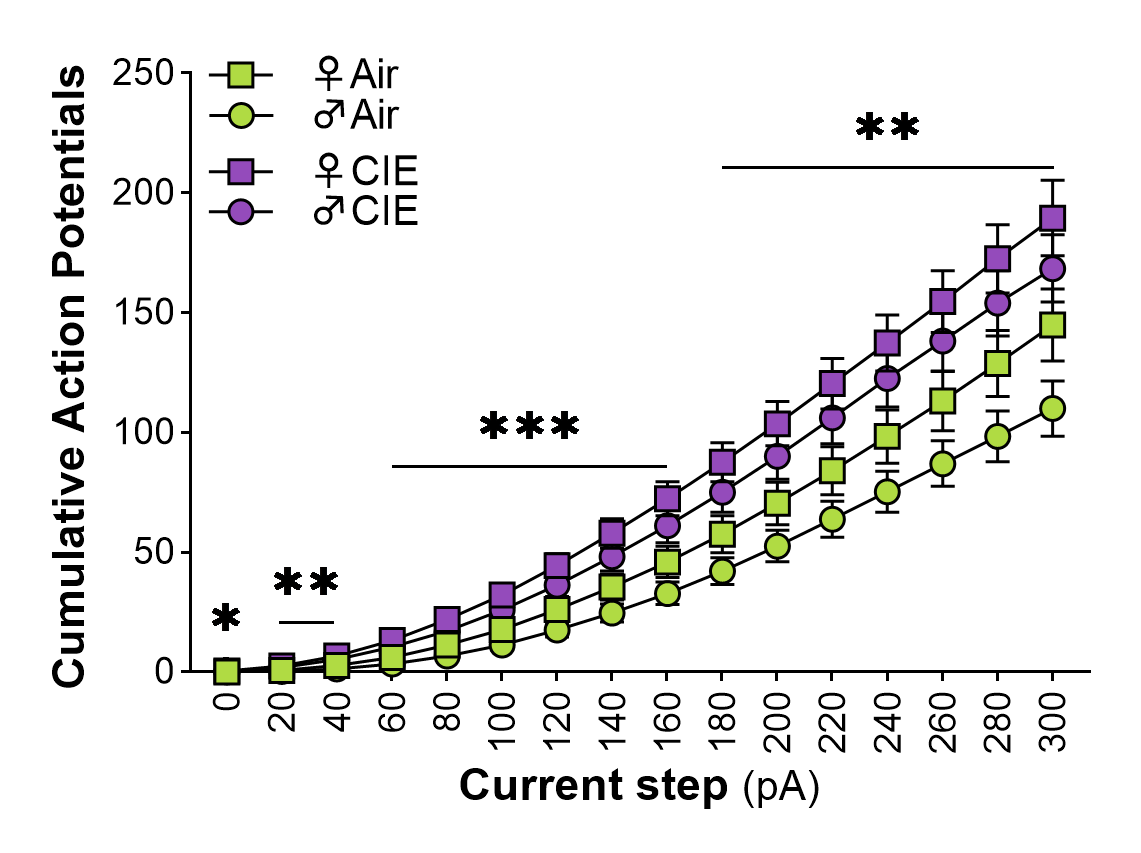

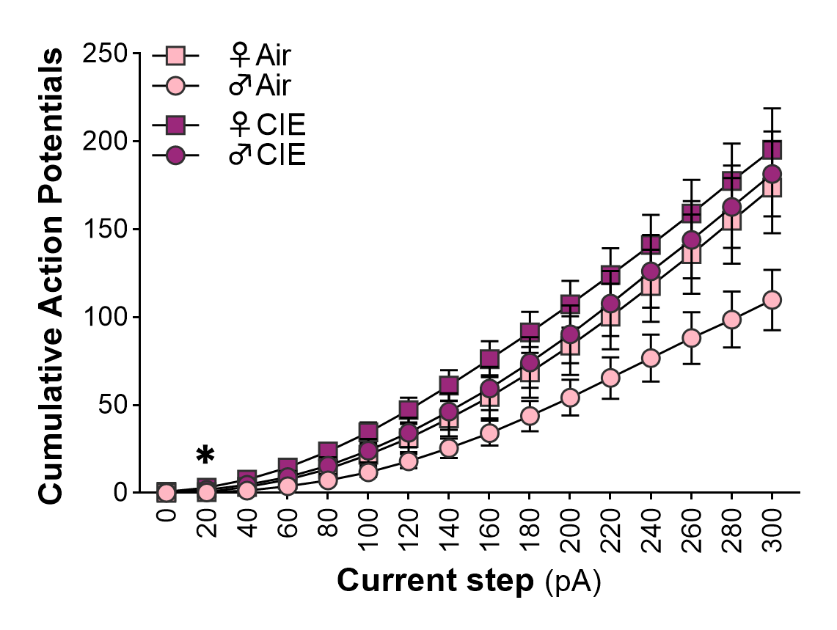

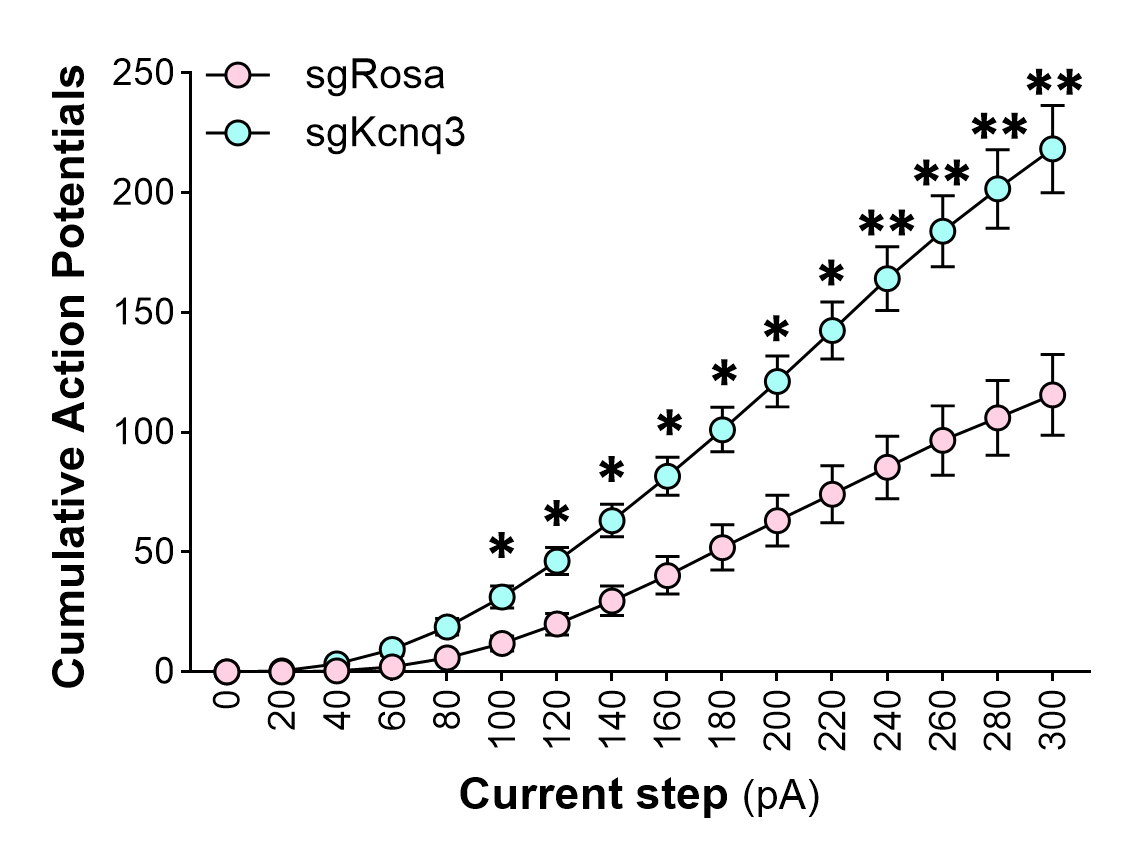

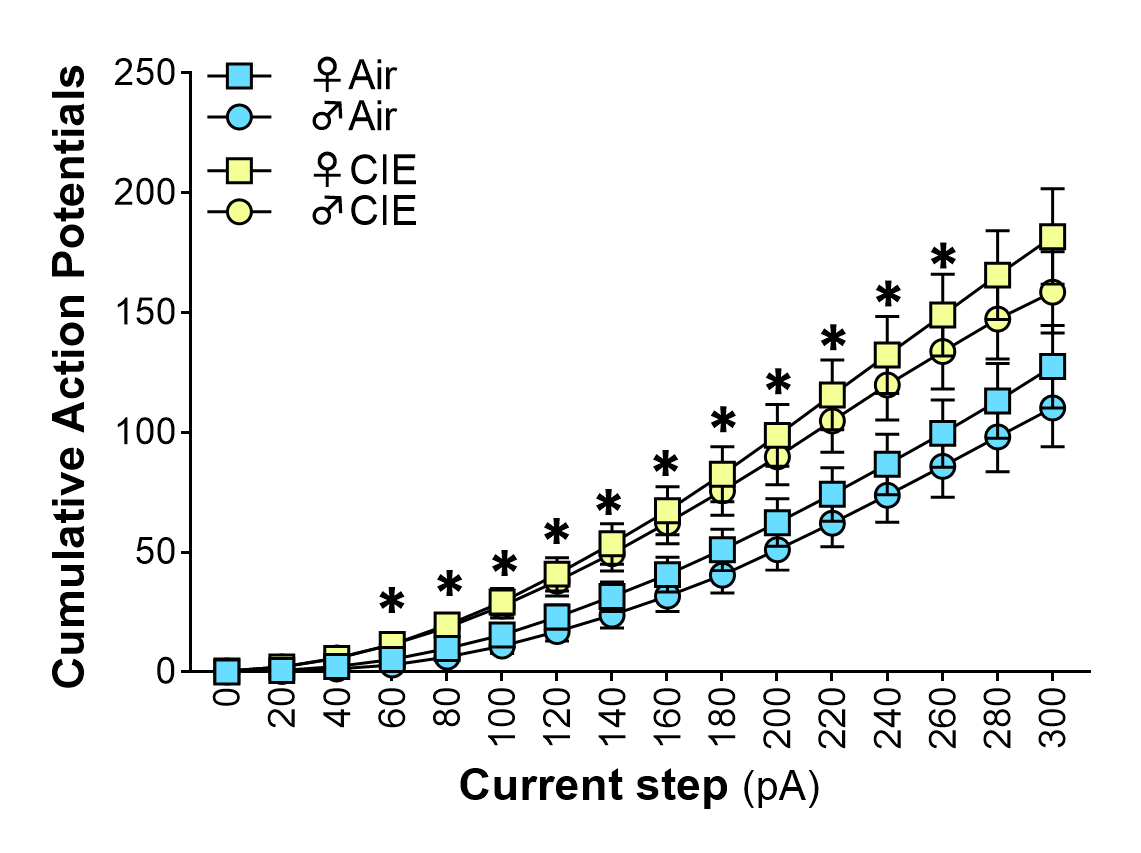

**B**

**A**

**D**

**C**

**Supplementary Figure 1. Cumulative spiking in dmPFC projection neurons.** (**A**) Cumulative action potential spiking in all dmPFC neurons (Step × Treatment interaction: *F*_1.079,110.1_ = 12.65, *p* = 0.0004). (**B**) Cumulative action potential spiking in ET projection neurons (Step × Treatment interaction: *F*_1.068,50.18_ = 4.396, *p* = 0.0387). (**C**) Cumulative action potential spiking in IT projection neurons (Step × Treatment interaction: *F*_1.093,55.73_ = 8.867, *p* = 0.0034). (**D**) Cumulative action potential spiking in dmPFC neurons expressing either sgRosa or sg*Kcnq3* (Step × AAV interaction: *F*_1.263,25.26_ = 14.31, *p* = 0.0004). post hocs, **p* < 0.05, ***p* < 0.01, ****p* < 0.001.
